# GAVIN - Gene-Aware Variant INterpretation for medical sequencing

**DOI:** 10.1101/072330

**Authors:** K. Joeri van der Velde, Eddy N. de Boer, Cleo C. van Diemen, Birgit Sikkema-Raddatz, Kristin M. Abbott, Alain Knopperts, Lude Franke, Rolf H. Sijmons, Tom J. de Koning, Cisca Wijmenga, Richard J. Sinke, Morris A. Swertz

## Abstract

Here, we present GAVIN, a new method that delivers accurate classification of variants for next-generation sequencing molecular diagnostics. It is based on gene-specific calibrations of allele frequencies (from the ExAC database), effect impact (using SnpEff) and estimated deleteriousness (CADD scores) for >3,000 genes. In a benchmark on 18 clinical gene sets, we achieved a sensitivity of 91.6%, with a specificity of 78.2%. This accuracy was unmatched by 12 other tools we tested. We provide GAVIN as an online MOLGENIS service to annotate VCF files, and as open source executable for use in bioinformatic pipelines. It can be found at http://molgenis.org/gavin.

## BACKGROUND

Only a few years ago, the high costs and technological challenges of whole exome and whole genome sequencing were limiting their application. Today, the practice of human genome sequencing has become routine even within the healthcare sector. This is leading to new and daunting challenges for clinical and laboratory geneticists[1]. Interpreting the thousands of variations observed in DNA and determining which are pathogenic and which are benign is still difficult and time-consuming, even when variants are prioritized by state-of-the-art *in silico* prediction tools and heuristic filters[2]. Using the current, largely manual, variant classification protocols, it is not feasible to assess the thousands of genomes per year now produced in a single hospital. It is the challenge of variant assessment which now impedes the effective uptake of next-generation sequencing into routine medical practice.

The recently introduced CADD[3] scores are a promising alternative[4]. These are calculated on the output of multiple *in silico* tools in combination with other genomic features. They trained a computer model on variants that have either been under long-term selective evolutionary pressure or none at all. The result was an estimation of deleteriousness for variants in the human genome, whether already observed or not. It has been shown to be a strong and versatile predictor for pathogenicity[3]. These scores may be used to define a classifier that labels a variant with a CADD score of >15 as probably pathogenic and <15 as benign, as suggested by the CADD authors[5]. Unfortunately, clinicians and laboratories cannot rely on this single threshold approach. We have shown that individual genes differ in their cut-off thresholds for what should be considered the optimal boundary between pathogenic or benign[4]. This issue has been partly addressed by MSC[6] (Mutation Significance Cutoff), which provides gene-based CADD cut-off values to remove inconsequential variants safely from sequencing data. While MSC aims to quickly and reliably reduce the number of benign variants left to interpret, it was not developed to detect/classify pathogenic variants.

The challenge is thus to find robust algorithms that classify both pathogenic and benign variants accurately and that fit into existing best practice, diagnostic filtering protocols[7]. Implementing such tools is not trivial because genes have different levels of tolerance to various classes of variants that may be considered harmful[8]. In addition, the pathogenicity estimates for benign variants are intrinsically lower because these are more common and of less severe consequence on protein transcription. Comparing the prediction score distributions of pathogenic variants with those of typical benign variants is therefore biased and questionable. Using such an approach means it will be unclear how well a predictor truly performs if a benign variant shares many properties with known pathogenic variants. Here, we present GAVIN (Gene-Aware Variant INterpretation), a new method that addresses these issues by gene-specific calibrations on closely matched sets of variants. GAVIN delivers accurate and reliable automated classification of variants for clinical application.

## RESULTS

### Development of GAVIN

GAVIN classifies variants as Benign, Pathogenic or a Variant of Uncertain Significance (VUS). It considers ExAC[8] minor allele frequency, SnpEff[9] impact and CADD score using gene-specific thresholds. For each gene, we ascertained ExAC allele frequencies and effect impact distributions of variants described in ClinVar (November 2015 release) [10] as pathogenic or likely pathogenic. From the same genes we selected ExAC variants that were not present in ClinVar as a benign reference set. We stratified this benign set to match the pathogenic set with respect to the effect impact distribution and minor allele frequencies (MAF). Using these comparable variant sets we calculated gene-specific mean values for CADD scores and minor allele frequencies as well as 95^th^ percentile sensitivity/specificity thresholds for both benign and pathogenic variants. We used fixed genome-wide classification thresholds as a fall-back strategy based on CADD scores < 15 for benign and > 15 for pathogenic[5] and on a MAF threshold of > 0.00474, which was the mean of all gene-specific pathogenic 95^th^ percentile thresholds. This allowed classification when insufficient variant training data were available to allow for gene-specific calibrations, or when the gene-specific rules failed to classify a variant. Based on the gene calibrations we then implemented GAVIN, which can be used online or via commandline (see http://molgenis.org/gavin) to perform variant classification.

### Performance benchmark

To test the robustness of GAVIN, we evaluated its performance using six benchmark variant classification sets from VariBench[11], MutationTaster2[12], ClinVar (only recently added variants that were not used for calibrating GAVIN), and a high-quality variant classification list from the University Medical Center Groningen (UMCG) genome diagnostics laboratory. These sets and the origins of their variants and classifications are described in Table 1. The combined set comprises 25,765 variants (17,063 benign, 8,702 pathogenic). All variants were annotated by SnpEff, ExAC and CADD prior to classification by GAVIN. To assess the clinical relevance of our method, we stratified the combined set into clinically relevant variant subsets based on organ-system specific genes. We formed 19 subset panels such as Cardiovascular, Dermatologic, and Oncologic based on the gene-associated physical manifestation categories from Clinical Genomics Database[13]. A total of 11,679 out of 25,765 variants were not linked to clinically characterized genes and formed a separate panel (see Table 2 for an overview). In addition, we assessed the performance of GAVIN in compared to 12 common *in silico* tools for pathogenicity prediction: MSC (using two different settings), CADD, SIFT[14], PolyPhen2[15], PROVEAN[16], Condel[17], PON-P2[18], PredictSNP2[19], FATHMM-MKL[20], GWAVA[21], FunSeq[22] and DANN[23].

**Table 1.** Variant and classification origins of the benchmark data sets used.

| Data set | Nr. of benign variants | Nr. of pathogenic variants | Origin |
| --- | --- | --- | --- |
| VariBench tolerance DS7, training set | 11,347 | 6,143 | PhenCode database, IDbases, and 18 individual LSDBs |
| VariBench tolerance DS7, test set | 1,377 | 510 | PhenCode database, IDbases, and 18 individual LSDBs |
| MutationTaster2 benchmark set | 1,194 | 161 | HGMD Professional and 1000 Genomes |
| ClinVar (additions of Nov 2015 to Feb 2016) | 1,668 | 1,688 | Submissions by clinical molecular geneticists, expert panels, diagnostic laboratories and companies |
| UMCG, variants exported from clinical diagnostic interpretation software | 1,176 | 174 | Clinical diagnostic classifications of variants in cardiology, dermatology, epilepsy, dystonia and preconception screening |
| UMCG, germline variants for familial cancer cases | 301 | 26 | Hereditary cancer variant classifications by an M.D. following ACMG guidelines |
| **Total** | **17,063** | **8,702** | **25,765** |

**Table 2.** Stratification of the combined variant data set into manifestation categories. The categories are defined by Clinical Genomics Database and are associated to clinically relevant genes. Variants were allocated to the manifestation categories based on their gene, and were placed in multiple categories if a gene was associated to multiple manifestations.

| **CGD manifestation panel** | **Genes** | **Variants** |
| --- | --- | --- |
| Allergy / Immunology / Infectious | 253 | 1,952 |
| Audiologic / Otolaryngologic | 217 | 1,215 |
| Biochemical | 354 | 2,538 |
| Cardiovascular | 446 | 4,360 |
| Craniofacial | 387 | 1,861 |
| Dental | 80 | 783 |
| Dermatologic | 345 | 2,749 |
| Endocrine | 240 | 1,801 |
| Gastrointestinal | 338 | 2,351 |
| Genitourinary | 149 | 1,026 |
| Hematologic | 267 | 2,571 |
| Musculoskeletal | 676 | 4,935 |
| Neurologic | 1,012 | 6,363 |
| Obstetric | 34 | 223 |
| Oncologic | 203 | 2,157 |
| Ophthalmologic | 479 | 3,649 |
| Pulmonary | 90 | 717 |
| Renal | 302 | 2,143 |
| *NotInCGD* | *5,806* | *11,679* |

Across all test sets, GAVIN achieved a median sensitivity of 91.6% and a median specificity of 78.2%. Other tools with >90% sensitivity were CADD (93.6%, specificity 57.1%) and MSC (97.1%, specificity 25.7%). The only other tool with >70% specificity was PredictSNP2 (70.6%, sensitivity 66.8%) (see Table 3 for an overview of tool performance). In all the clinical gene sets GAVIN scored >90% sensitivity, including >93% for Cardiovascular, Biochemical, Obstetric and Dermatologic genes. The non-clinical genes scored 71.3%. The specificity in clinical subsets ranged from 71.6% for Endocrine to 83.8% for Dental. Non-clinical gene variants were predicted at 70.2% specificity. See **Supplementary Table 1** for detailed results.

**Table 3.** Performance overview of all tested tools.

| **Tool** | **Median Sensitivity** | **Median Specificity** |
| --- | --- | --- |
| CADD | 93.6% | 57.1% |
| Condel | 70.3% | 39.5% |
| DANN | 63.8% | 66.7% |
| FATHMM | 69.5% | 61.9% |
| FunSeq | 61.7% | 50.2% |
| GAVIN | 91.6% | 78.2% |
| GWAVA | 47.6% | 26.2% |
| MSC_ClinVar95CI | 84.7% | 64.4% |
| MSC_HGMD99CI | 97.1% | 25.7% |
| PolyPhen2 | 68.0% | 46.8% |
| PONP2 | 47.5% | 26.9% |
| PredictSNP2 | 66.8% | 70.6% |
| PROVEAN | 65.9% | 62.1% |
| SIFT | 67.9% | 57.9% |

We illustrated the practical implications of classification sensitivity and specificity in Table 4. Here, 90%/80% represents the performance of GAVIN, 90%/60% matches CADD, and 70%/80% or 70%/60% can be considered averages of other methods. In a hypothetical example where 110 variants are being tested (100 benign and 10 pathogenic), the difference in predictive value between the performance opposites is over two-fold (31% positive predictive value (PPV) for 90/80% and 15% PPV for 70/60%).

**Table 4.**
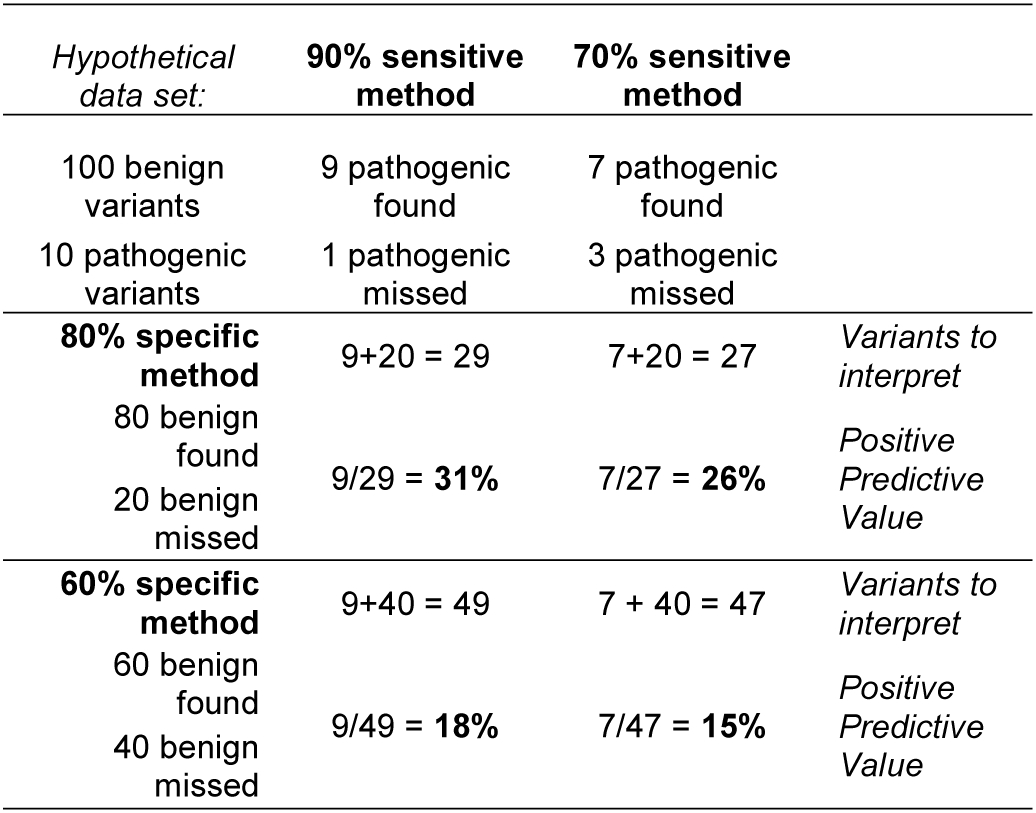
The practical impact in clinical diagnostics of using methods of different sensitivity and specificity on a data set with 100 benign and 10 pathogenic variants.

### Added value of gene-specific calibration

We then investigated the added value of using gene-specific thresholds on classification performance relative to using genome-wide thresholds. We bootstrapped the performance on 10,000 random samples of 100 benign and 100 pathogenic variants. These variants were drawn from the three groups of genes described in Materials & Methods: (1) genes for which CADD was significantly predictive for pathogenicity (n = 520), (2) genes where CADD was not significantly predictive (n = 660), and (3) genes with scarce variant data available for calibration (n = 737). For each of these sets we compared the use of gene-specific CADD and MAF classification thresholds with that of genome-wide filtering rules (CADD score < 15 and MAF >0.00474 for benign, otherwise classify as pathogenic).

We observed the highest accuracy on genes for which CADD had significant predictive value and for the gene-specific classification method (median accuracy = 87.5%); this was significantly higher than using the genome-wide method for these same genes (median accuracy = 85%, Mann-Whitney U test p-value < 2.2e-16). For genes for which CADD had less predictive value we found a lower overall performance, but still reached a significantly better result using the gene-specific approach (median accuracy = 84.5% versus genome-wide 82%, p-value < 2.2e-16). Lastly, the worst performance was seen for variants in genes with scarce training data available. The gene-specific performance, however, was still significantly better than using genome-wide thresholds (median accuracy = 83.5% and 81% respectively, p-value = 2.2e-16). See Figure 2.

**Figure 1.**
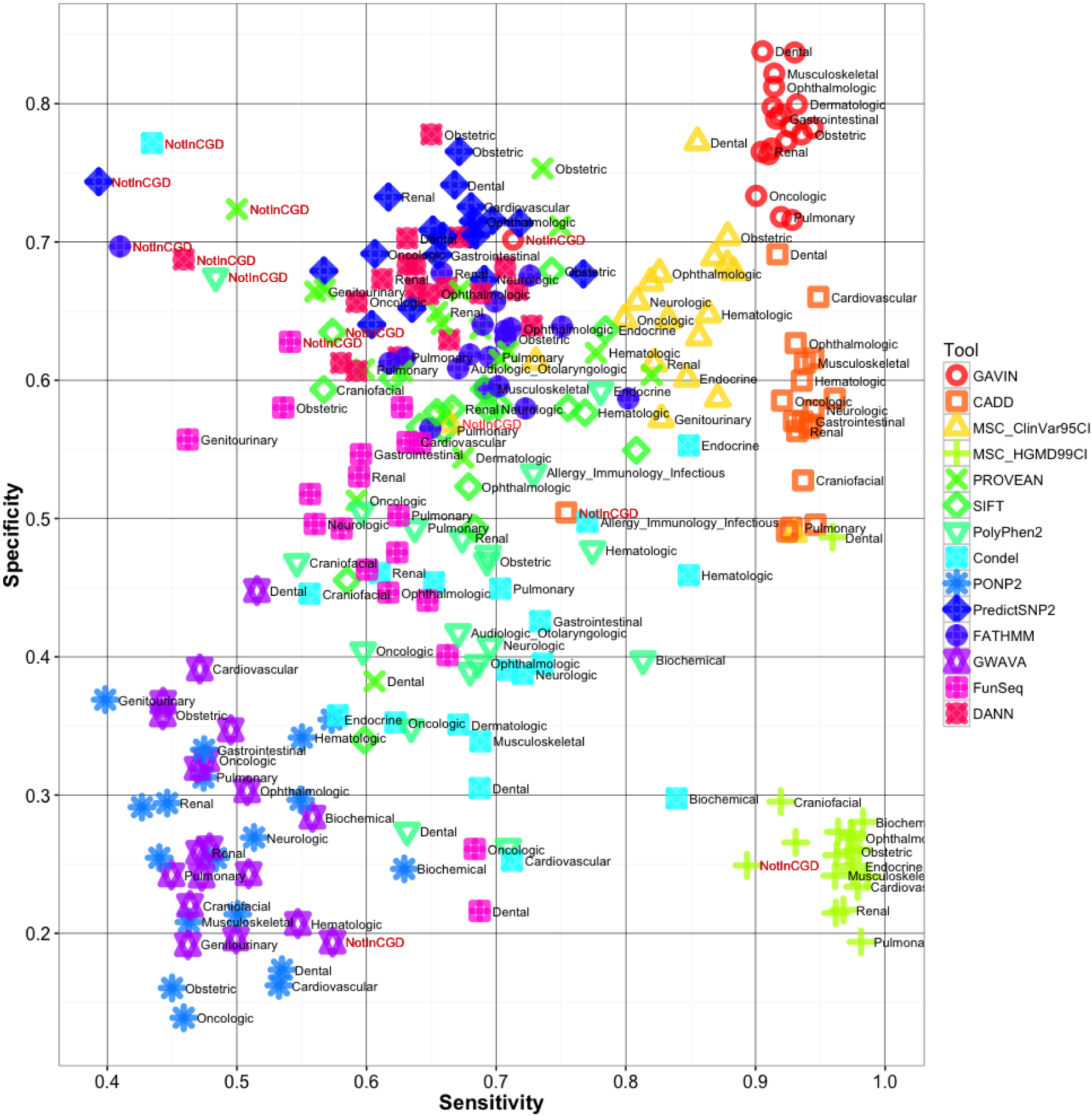
Performance of GAVIN and other tools across different clinical gene sets. Prediction 537 quality is measured as sensitivity and specificity, i.e. the fraction of pathogenic variants correctly 538 identified and the fraction of mistakes made while doing so.

**Figure 2.**
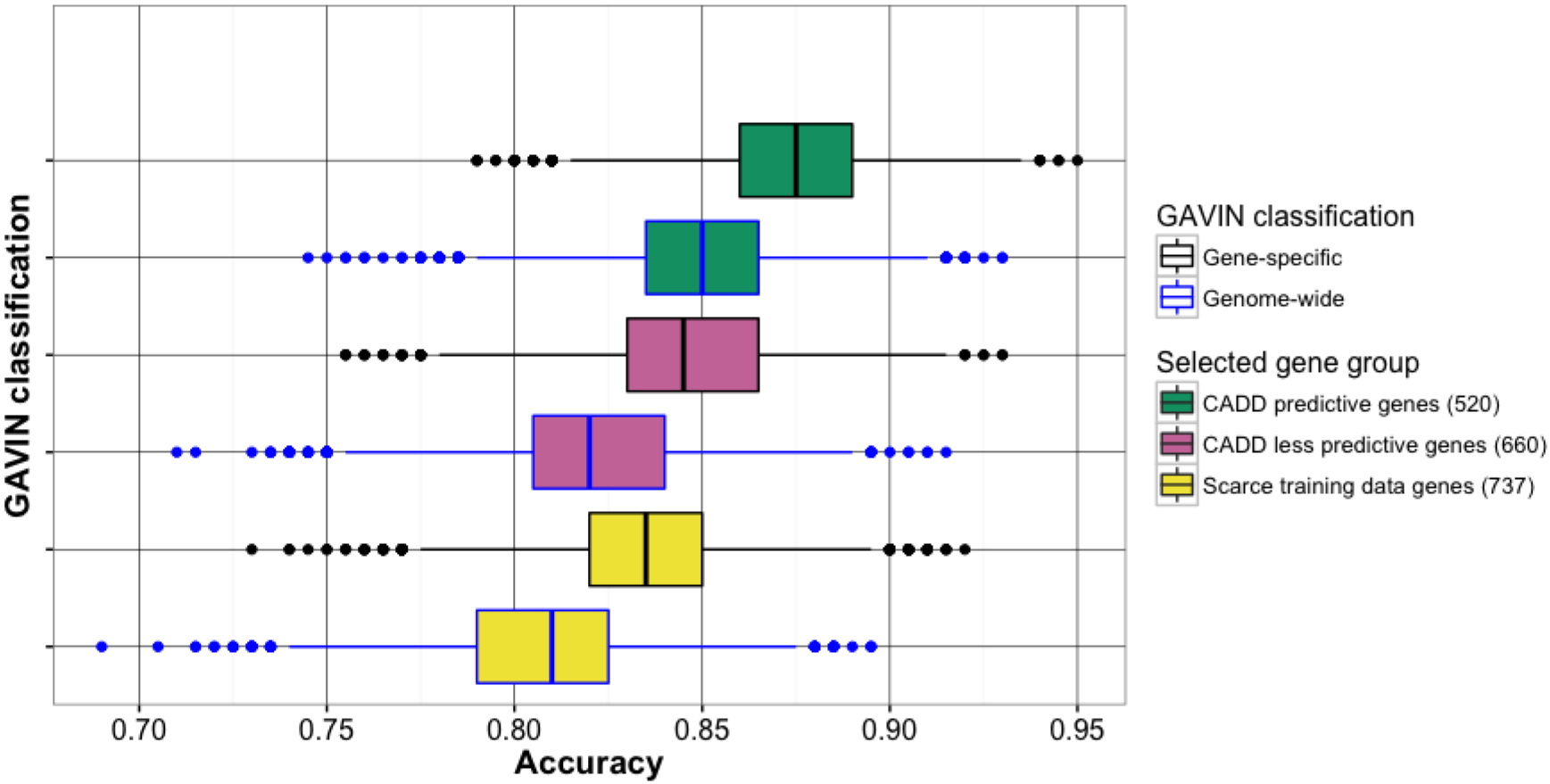
Comparison of gene-specific classification thresholds with genome-wide fixed thresholds in three groups of genes: 520 genes for which CADD is predictive, 660 genes for which CADD is less predictive, and 737 genes with scarce training data. For each group, 10,000 sets of 100 benign and 100 pathogenic variants were randomly sampled and tested from the full set of 25,765 variants and accuracy was calculated for gene-specific and genome-wide CADD and MAF thresholds.

## DISCUSSION

We have developed GAVIN, a method for automated variant classification using gene-specific calibration of classification thresholds for benign and pathogenic variants.

Our results show that GAVIN is a powerful classifier with consistently high performance in clinically relevant genes. The robustness of our method arises from a calibration strategy that first corrects for calibration bias between benign and pathogenic variants, in terms of consequence and rarity, before calculating the classification thresholds. A comprehensive benchmark demonstrates a unique combination of high sensitivity (>90%) and high specificity (>70%) for variants in genes related to different organ systems. This is a significant improvement over existing tools that tend to achieve either a high sensitivity (CADD, MSC) or a high specificity (PredictSNP2). A high sensitivity is crucial for clinical interpretation because pathogenic variants should not be falsely discarded. In addition, having a higher specificity means that the results will be far less ‘polluted’ with false-positives and thus less risk of patients being given a wrong molecular diagnosis. GAVIN decreases false-positives by about 20% compared to using CADD for the same purpose, thereby reducing the interpretation time considerably. The difference between using a high and low performance method can be dramatic in practice. In a hypothetical example, GAVIN would make downstream variant interpretation twice as effective as a low performance method, with more sensitive detection of pathogenic variants.

Even though an optimal combination of sensitivity and specificity may be favorable in general terms, there may still be a need for tools that perform differently. The MSC gene-specific thresholds based on HGMD[24] at 99% confidence interval show a very high sensitivity (97.1%), but at the expense of a very low specificity (25.7%). Such low specificity thresholds will pick up almost all the pathogenic variants with scores exceeding gene thresholds. This allows safe removal (<3% error) of benign variants that fall below these thresholds, which was their authors’ aim. However, this tool cannot detect pathogenic variants due its low specificity. Other tools, such as PON-P2, may show a relatively low performance, but not necessarily because of true errors. Such tools may simply be very ‘picky’ and only return a classification when the verdict carries high confidence. If we ignore the variants that PON-P2 did not classify, and only consider how many of the variants that it did classify were correct, we find a positive predictive value of 96%, and a negative predictive value of 94%. Thus, while this tool might not be useful for exome screening because too many pathogenic variants would be lost, it can still be an excellent choice for further investigation of interesting variants. We would therefore emphasize that appropriate tools should be selected depending on the question or analysis protocol used and by taking their strengths and weaknesses into account.

Not surprisingly, we could confirm that the use of gene-specific thresholds instead of genome-wide thresholds led to a consistent and significant improvement of classification performance. This shows the added value of our strategy. Overall performance was slightly lower in genes for which CADD has limited predictive value, and even lower in genes with few ‘gold standard’ pathogenicity data available. Evaluating variants in uncharacterized genes is rare in clinical diagnostics, although it may occur when exome sequencing is aimed at solving complex phenotypes or undiagnosed cases. Nevertheless, GAVIN is likely to improve continuously in an increasing number of genes, propelled by the speed at which pathogenic variants are now being reported.

With GAVIN we were also able to demonstrate the residual power of CADD scores as a predictor for pathogenicity on a gene-by-gene basis, revealing that the scores are informative for many genes (these results can be accessed at http://molgenis.org/gavin). There are several possible explanations for potential non-informativity of CADD scores. It may have bias towards the *in silico* tools and sources it was trained on, limiting their predictiveness for certain genomic regions or disease mechanisms[25]. Furthermore, calibration of pathogenic variants could be difficult in genes with high damage tolerance, i.e. having many missense or loss-of-function mutations[26]. In addition, calibration may be impaired by false input signals, such as an incorrect pathogenic classification in ClinVar or inclusion of disease cohorts in large databases such as ExAC could misrepresent allele frequencies[27]. Lastly, pathogenic variants could have a low penetrance or their effect mitigated by genetic modifiers, causing high deleteriousness to be tolerated in the general population against expectations[28].

The field of clinical genomics is now moving towards interpretation of non-coding disease variants (NCVs) identified by whole-genome sequencing[29]. A number of recently introduced metrics, including EIGEN[30], FATHMM-MKL, DeepSEA[31], and GWAVA, specialize in predicting the functional effects of non-coding sequence variation. When a pathogenic NCV reference set of reasonable quantity becomes available, a calibration strategy as described here will be essential to be able to use these metrics effectively in whole-genome diagnostics.

## CONCLUSIONS

GAVIN provides an automated decision-support protocol for classifying variants, which will continue to improve in scope and precision as more data is publicly shared by genome diagnostic laboratories. Our approach bridges the gap between estimates of genome-wide and population-wide variant pathogenicity and contributes to their practical usefulness for interpreting clinical variants in specific patient populations. Databases such as ClinVar contain a wealth of implicit rules now used manually by human experts to classify variants. These rules are deduced and employed by GAVIN to classify variants that have not been seen before.

We envision GAVIN accelerating NGS diagnostics and becoming particularly beneficial as a powerful (clinical) exome screening tool. It can be used to quickly and effectively detect over 90% of pathogenic variants in a given data set and to present these results with an unprecedented small number of false-positives. It may especially serve laboratories that lack the resources necessary to perform reliable and large-scale manual variant interpretation for their patients, and spur the development of more advanced gene-specific classification methods. We provide GAVIN as an online MOLGENIS[32] web service to browse gene calibration results and annotate VCF files, and as a commandline executable including open source code for use in bioinformatic pipelines. GAVIN can be found at http://molgenis.org/gavin.

## METHODS

### Calibration of gene-specific thresholds

We downloaded ClinVar (variant_summary.txt.gz from ClinVar FTP, last modified date: 05/11/15) and selected GRCh37 variants that contained the word “pathogenic” in their clinical significance. These variants were matched against the ClinVar VCF release (clinvar.vcf.gz, last modified date: 01/10/15) using RS (Reference SNP) identifiers in order to resolve missing indel notations. On the resulting VCF, we ran SnpEff version 4.1L with these settings: hg19 -noStats -noLog -lof -canon -ud 0. As a benign reference set, we selected variants from ExAC (release 0.3, all sites) from the same genic regions with +/− 100 bases of padding on each side to capture more variants residing on the same exon.

We first determined the thresholds for gene-specific pathogenic allele frequency by taking the ExAC allele frequency of each pathogenic variant, or assigning zero if the variant was not present in ExAC, and calculating the 95^th^ percentile value per gene using the R7 method from Apache Commons Math version 3.5. We filtered the set of benign variants with this threshold to retain only variants that were rare enough to fall into the pathogenic frequency range.

Following this step, the pathogenic impact distribution was calculated as the relative proportion of the generalized effect impact categories, as annotated by SnpEff on the pathogenic variants. The same calculation was performed for the benign variants using the variant Ensembl VEP[33] consequence types already present in ExAC. To facilitate this, we defined a trivial mapping of VEP consequences types (being equivalent to SnpEff consequences) to SnpEff impact categories. The benign variants were subsequently downsized to match the impact distribution of the pathogenic variants.

For instance, in the case of 407 pathogenic MYH7 variants, we found a pathogenic allele frequency threshold of 9.494e-05, and an impact distribution of 5.41% HIGH, 77.4% MODERATE, 17.2% LOW and 0% MODIFIER. We defined a matching set of benign variants by retrieving 1,799 MYH7 variants from ExAC (impact distribution: 2.1% HIGH, 23.52% MODERATE, 32.07% LOW, 42.32% MODIFIER), from which we excluded known ClinVar pathogenic variants (n = 99), variants above the AF threshold (n = 377), and removed interspersed variants using a non-random ‘step over’ algorithm until the impact distribution was equalized (n = 862). We thus reached an equalized set of 461 variants. This process was repeated for 3,055 genes.

We then obtained the CADD scores for all variants and tested whether there was a significant difference in scores between the sets of pathogenic and benign variants for each gene, using a Mann-Whitney U test. Per gene we determined the mean CADD score for each group, and also the 95^th^ percentile sensitivity threshold (detection of most pathogenic variants while accepting false-positives) and 95^th^ percentile specificity threshold (detection of most benign variants while accepting false-negatives), using the Percentile R7 function. All statistics were done with Apache Commons Math version 3.5.

On average, CADD scores were informative of pathogenicity. The mean benign variant CADD score across all genes was 23.68, while the mean pathogenic variant CADD score was 28.45, a mean difference of 4.77 (σ = 4.69). Of 3,055 genes that underwent the calibration process, we found 520 “CADD predictive” genes that had a significantly higher CADD score for pathogenic variants than for benign variants (Mann-Whitney U test, p-value <0.05). Interestingly, we also found 660 “CADD less predictive” genes, for which there was no proven difference between benign and pathogenic variants (p-value >0.05 despite having ≥ 5 pathogenic and ≥5 benign variants in the gene). For 737 genes there was very little calibration data available (<5 pathogenic or <5 benign variants), resulting in no significant difference (p-value >0.05) between CADD scores of pathogenic and benign variants. We found 309 genes for which effect impact alone was predictive, meaning that a certain impact category was unique for pathogenic variants compared to benign variants. For instance, when observing HIGH impact pathogenic variants (frame shift, stopgain, etc.) for a given gene, whereas benign variants only reached MODERATE impact (missense, inframe insertion, etc.). No further CADD calibration was performed on these genes. See http://www.molgenis.org/gavin for a full table of gene calibration outcomes.

### Variant sets for benchmarking

We obtained six variant sets that had been classified by human experts. These data sets were used to benchmark the *in silico* variant pathogenicity prediction tools mentioned in this paper. Variants from the original sets may sometimes be lost due to conversion of cDNA/HGVS notation to VCF.

The VariBench protein tolerance data set 7 (http://structure.bmc.lu.se/VariBench/) contains disease-causing missense variations from the PhenCode[34] database, IDbases[35], and 18 individual LSDBs[11]. The training set we used contained 17,490 variants, of which 11,347 were benign and 6,143 pathogenic. The test set contained 1,887 variants, of which 1,377 were benign and 510 pathogenic. We used both the training set and test set as benchmarking sets.

The MutationTaster2[12] test set contains known disease mutations from HGMD[24] Professional and putatively harmless polymorphisms from 1000 Genomes. It is available at http://www.mutationtaster.org/info/Comparison_20130328_with_results_ClinVar.html. This set contains 1,355 variants, of which 1,194 are benign and 161 pathogenic.

We selected 1,688 pathogenic variants from ClinVar that were added between November 2015 and February 2016 as an additional benchmarking set, since our method was based on the November 2015 release of ClinVar. We supplemented this set with a random selection of 1,668 benign variants from ClinVar, yielding a total of 3,356 variants.

We obtained an in-house list of 2,359 variants that had been classified by molecular and clinical geneticists at the University Medical Center Groningen. These variants belong to patients seen in the context of various disorders: cardiomyopathies, epilepsy, dystonia, preconception carrier screening, and dermatology. Variants were analyzed according to Dutch medical center guidelines[36] for variant interpretation, using Cartagenia Bench Lab^TM^ (Agilent Technologies) and Alamut® software (Interactive Biosoftware) by evaluating in-house databases, known population databases (1000G[37], ExAC, ESP6500 at http://evs.gs.washington.edu/EVS/, GoNL[38]), functional effect and literature searches. Any ClinVar variants included in the November 2015 release were removed from this set to prevent circular reasoning, resulting in a total of 1,512 variants, with 1,176 benign/likely benign (merged as Benign), 162 VUS, and 174 pathogenic/likely pathogenic (merged as Pathogenic).

From the UMCG diagnostics laboratory we also obtained a list of 607 variants seen in the context of familial cancers. These were interpreted by a medical doctor according to ACMG guidelines[7]. We removed any ClinVar variants (November 2015 release), resulting in 395 variants, with 301 benign/likely benign (merged as Benign), 68 VUS and 26 likely pathogenic/pathogenic (merged as Pathogenic).

### Variant data processing and preparation

We used Ensembl VEP (http://grch37.ensembl.org/Homo_sapiens/Tools/VEP/) to convert cDNA/HGVS notations to VCF format. Newly introduced N-notated reference bases were replaced with the appropriate GRCh37 base, and alleles were trimmed where needed (e.g. “TA/TTA” to “T/TT”). We annotated with SnpEff (version 4.2) using the following settings: hg19 -noStats -noLog -lof -canon -ud 0. CADD scores (version 1.3) were added by running the variants through the CADD webservice (available at http://cadd.gs.washington.edu/score). ExAC (release 0.3) allele frequencies were added with MOLGENIS annotator (release 1.16.2). We also merged all benchmarking sets into a combined file with 25,995 variants (of which 25,765 classified as benign, likely benign, likely pathogenic or pathogenic) for submission to various online *in silico* prediction tools.

### Execution of *in silico* predictors

The combined set of 25,765 variants was classified by the *in silico* variant pathogenicity predictors (MSC, CADD, SIFT, PolyPhen2, PROVEAN, Condel, PON-P2, PredictSNP2, FATHMM, GWAVA, FunSeq, DANN). The output of each tool was loaded into a program that compared the observed output to the expected classification and which then calculated performance metrics such as sensitivity and specificity. The tools that we evaluated and the web addresses used can be found in **Supplementary Table 2**. We executed PROVEAN and SIFT, for which the output was reduced by retaining the following columns: “INPUT”, “PROVEAN PREDICTION (cut-off = −2.5)” and “SIFT PREDICTION (cut-off = 0.05)”. For PONP-2, the output was left as-is. The Mutation Significance Cutoff (MSC) thresholds are configurable; we downloaded the ClinVar-based thresholds for CADD 1.3 at 95% confidence interval, comparable to our method, as well as HGMD-based thresholds at 99% confidence interval, the default setting. Variants below the gene-specific thresholds were considered benign, and above the threshold pathogenic. We obtained CADD scores of version 1.3. Following the suggestion of the CADD authors, scores of variants below a threshold of 15 were considered benign, above this threshold pathogenic. The output of Condel was reduced by retaining the following columns: “CHR”, “START”, “SYMBOL”, “REF”, “ALT”, “MA”, “FATHMM”, “CONDEL”, “CONDEL_LABEL”. After running PolyPhen2, its output was reduced by retaining the positional information (“chr2:220285283|CG”) and the “prediction” column. Finally, we executed PredictSNP2, which contains the output from multiple tools. From the output VCF, we used the INFO fields “PSNPE”, “FATE”, “GWAVAE”, “DANNE” and “FUNE” for the pathogenicity estimation outcomes according to the PredictSNP protocol for PredictSNP2 consensus, FATHMM, GWAVA, DANN and FunSeq, respectively.

### Stratification of variants using Clinical Genomics Database

We downloaded Clinical Genomics Database (CGD; the .tsv.gz version on 1 June 2016 from http://research.nhgri.nih.gov/CGD/download/). A Java program evaluated each variant in the full set of 25,765 variants and retrieved their associate gene symbols as annotated by SnpEff. We matched the gene symbols to the genes present in CGD and retrieved the corresponding physical manifestation categories. Variants were then written out to separate files for each manifestation category (cardiovascular, craniofacial, renal, etc.). This means a variant may be output into multiple files if its gene was linked to multiple manifestation categories. However, we did prevent variants from being written out twice to the same file in the case of overlapping genes in the same manifestation categories. We output a variant into the “NotInCGD” file only if it was not located in any gene present in CGD.

### Implementation

GAVIN was implemented using Java 1.8 and MOLGENIS[32] 1.16 (http://molgenis.org). Source code with tool implementation details can be found at https://github.com/molgenis/gavin. All benchmarking and bootstrapping tools, as well as all data processing and calibration tools, can also be found in this source code repository.

### Binary classification metrics

Prediction tools may classify variants as benign or pathogenic, but may also fail to reach a classification or classify a variant as VUS. Because of these three outcome states, binary classification metrics must be used with caution. According to standard definitions of ‘sensitivity’, such as the following example: “Recall or Sensitivity (as it is called in Psychology) is the proportion of Real Positive cases that are correctly Predicted Positive” (source: https://csem.flinders.edu.au/research/techreps/SIE07001.pdf), we define sensitivity as the number of detected pathogenic variants (true-positives) over the total number of pathogenic variants, which includes true-positives, false-negatives (pathogenic variants misclassified as benign), and pathogenic variants that were otherwise ‘missed’, i.e. classified as VUS or not classified at all. Therefore, Sensitivity = TruePositive/(TruePositive + False-Negative + MissedPositive). We applied the same definition for specificity, and define it as: Specificity = TrueNegative/(TrueNegative + FalsePositive + MissedNegative). Following this line, accuracy is then defined as (TP + TN)/(TP + TN + FP + FN + MissedPositive + MissedNegative).

## DECLARATIONS

### Ethics approval and consent to participate

The study was done in accordance with the regulations and ethical guidelines of the University Medical Center Groningen. Specific ethical approval was not necessary because this study was conducted on aggregated, fully anonymized data.

### Consent for publication

Not applicable.

### Availability of data and material

The datasets generated during and/or analysed during the current study are available in the GAVIN public GitHub repository, available at https://github.com/molgenis/gavin.

### Authors' contributions

KV, EB, MS conceived the method. KV, EB, CD, BS, KA, LF, CW, RHS, RJS and TK helped to fine-tune the method, accumulate relevant validation data and evaluate the results. KV, MS drafted the manuscript. KV, EB, CD, BS, KA, AK, LF, FS, TK, CW, RHS, RJS, MS edited and reviewed the manuscript.

## Acknowledgements

We thank Jackie Senior, Kate Mc Intyre and Diane Black for editorial advice. We thank the MOLGENIS team for assistance with the software implementation and the GAVIN user interface: Bart Charbon, Fleur Kelpin, Mark de Haan, Erwin Winder, Tommy de Boer, Jonathan Jetten, Dennis Hendriksen, Chao Pang.

---

**Supplementary Table 1.**

Detailed overview of all benchmark results. Each combination of tool and data set is listed. We provide the raw counts of true-positives (TP), true-negatives (TN), false-positives (FP) and false-negatives (FN), as well as of pathogenic and benign variants that were ‘missed’, i.e. not correctly identified as such. From these numbers we calculated the sensitivity and specificity.

**Supplementary Table 2.**

The tools used to evaluate our benchmark variant set, and the web addresses used through which they were accessed.

